# Micromagnetic Stimulation (μMS) Dose-Response of the Rat Sciatic Nerve

**DOI:** 10.1101/2022.11.23.517726

**Authors:** Renata Saha, Zachary Sanger, Robert Bloom, Onri J. Benally, Kai Wu, Denis Tonini, Walter C. Low, Susan A. Keirstead, Theoden I. Netoff, Jian-Ping Wang

## Abstract

**Objective:** The objective of this study was to investigate the effects of micromagnetic stimuli strength and frequency from the <u>Mag</u>netic <u>Pen</u> (MagPen) on the rat right sciatic nerve. The nerve’s response would be measured by recording muscle activity and movement of the right hind limb.

**Approach:** The MagPen was custom-built such that it can be held over the sciatic nerve in a stable manner. Rat leg muscle twitches were captured on video and movements were extracted using image processing algorithms. EMG recordings were also used to measure muscle activity.

**Main results:** The MagPen prototype when driven by alternating current, generates time-varying magnetic field which as per Faraday’s Law of Electromagnetic Induction, induces an electric field for neuromodulation. The orientation dependent spatial contour maps for the induced electric field from the MagPen prototype has been numerically simulated. Furthermore, in this *in vivo* work on μMS, a dose-response relationship has been reported by experimentally studying how the varying amplitude (Range: 25 mV_p-p_ through 6 V_p-p_) and frequency (Range: 100 Hz through 5 kHz) of the MagPen stimuli alters the hind limb movement. The primary highlight of this dose-response relationship is that at a higher frequency of the μMS stimuli, significantly smaller amplitudes can trigger hind limb muscle twitch. This frequency-dependent activation can be justified following directly from the Faraday’s Law as the magnitude of the induced electric field is directly proportional to frequency.

**Significance:** This work reports that μMS can successfully activate the sciatic nerve in a dose-dependent manner. The MagPen probe, unlike electrodes, does not have a direct electrochemical interface with tissues rendering it much safer than an electrode. Magnetic fields create more precise activation than electrodes because they induce smaller volumes of activation. Finally, unique features of μMS such as orientation dependence, directionality and spatial selectivity have been demonstrated.

## 1. Introduction

Implantable neuromodulation devices have been used for treatment of chronic pain [1–3], brain disorders [4–7] as well as retinal [8,9] and cochlear prosthesis [8,9]. Using electric shock from Torpedo fish for treatment of chronic pain dates back to 15^th^ century AD [10] followed by reports of muscle contraction due to electrical stimulation [11,12] in the 18^th^ century. However, the modern era of neuromodulation for treatment of chronic pain began in the 1960s [13]. *Sciatica* is a diagnostic condition where excruciating pain travels through the path of the sciatic nerve – the nerve originates from the spinal cord and extending through the legs. Transcutaneous electric nerve stimulation (TENS) [14,15] has been proven to be quite effective for *sciatica* treatment, and percutaneous electric nerve stimulation (PENS) is being investigated as a viable and possibly an even better alternative [16,17].

Electrical stimulation has been proven to be clinically safe and has a long and successful track record. However, it has several drawbacks which are yet to be addressed. In electrical stimulation, the electrodes are in close galvanic contact with biological tissue, which can result in biofouling [18,19]. Biofouling increases the necessary voltage required to achieve the same field strengths in the tissue and can cause field strengths necessary to activate the tissue beyond safe limits. Here we present a novel implantable technology based on micromagnetic stimulation (μMS). It uses submillimeter-sized coils, or microcoils (μcoils) that, when driven by a time-varying current, generate a time-varying magnetic field that induces an electric field in the surrounding tissue as per Faraday’s Laws of Electromagnetic Induction. Because stimulation is through an inductive effect, the μcoils do not come in direct electrochemical contact with the biological tissue. Therefore, these μcoils may not be as sensitive to biofouling as electrodes. Furthermore, μcoils may have advantages in MRI compatibility [20], spatial selectivity, and directionality [21]. The optimized shape of the coil for neuromodulation is still under investigation, with several designs reported so far both using computational modeling as well as experimental approaches [23–28]. Previous studies have demonstrated the efficacy of magnetic stimulation using cm-sized and mm-sized coils on the rat sciatic nerve [22], here we demonstrate stimulation using micromagnetic stimulation (μMS).

In this work, we present μMS using the <u>Mag</u>netic <u>Pen</u> (MagPen) [21] on the rat sciatic nerve. We measure a dose-response relationship of magnetic stimuli (varying amplitudes and durations) on twitch in the hind limb evoked via the sciatic nerve. In addition, we have studied the importance of directionality of the induced electric field from the μcoil w.r.t the nerve, the significance of angular orientation of the μcoil w.r.t the plane of the neural tissue and reported frequency dependent micromagnetic activation of the sciatic nerve with support from experimental observation. Overall, this work demonstrates the efficacy and feasibility of neuromodulation using μMS which may open a new scope for the study of *sciatica* and chronic pain management.

## 2. Materials and methods

### 2.1. MagPen: The micromagnetic neurostimulation probe fabrication

The fabrication for the micromagnetic neurostimulation probe, MagPen (see Fig. 1(a)) has been reported previously in [21] and the only difference is in the model of the μcoil. Regarding appropriate choice of the μcoil model, it was made sure that the electrical characteristics of the μcoil were reduced to that of a simple resistive-inductive (RL) circuit (see Table S1 for LCR meter measurements of the RLC characteristics). In this work, we used a commercially available μcoil (Model no.: TDK Corporation MLG1005SR10JTD25) which was soldered (using solder flux & hot air blower) at the tip of printed circuit boards (PCB) of length 3 cm. For ease of orientation of the MagPen over the nerve, we trimmed the thickness of the PCB down along the z axis to 0.4 mm (see Fig. 1(a)). This being the first report that stimulates the rat sciatic nerve, the orientation of the μcoil that would potentially activate the nerve was unknown; in fact, hard to predict. Hence, similar to our previous work [21], we fabricated the MagPen in two orientations: Type <u>H</u>orizontal (Type H) and Type <u>V</u>ertical (Type V) (see Fig. 1(a)). The tip of the MagPen which holds the μcoil has a width of 1.7 mm and 1.4 mm for the Type H and Type V respectively. The complete image of the prototype has been reported in Fig. S1 of Supplementary Information S1. The μcoil is of dimensions 1 mm × 600 μm × 500 μm and the tiny size of the μcoil is compared to the tip of a pencil (Dixon No.2 HB pencil) (see Fig. 1(b)).

**Figure 1.**
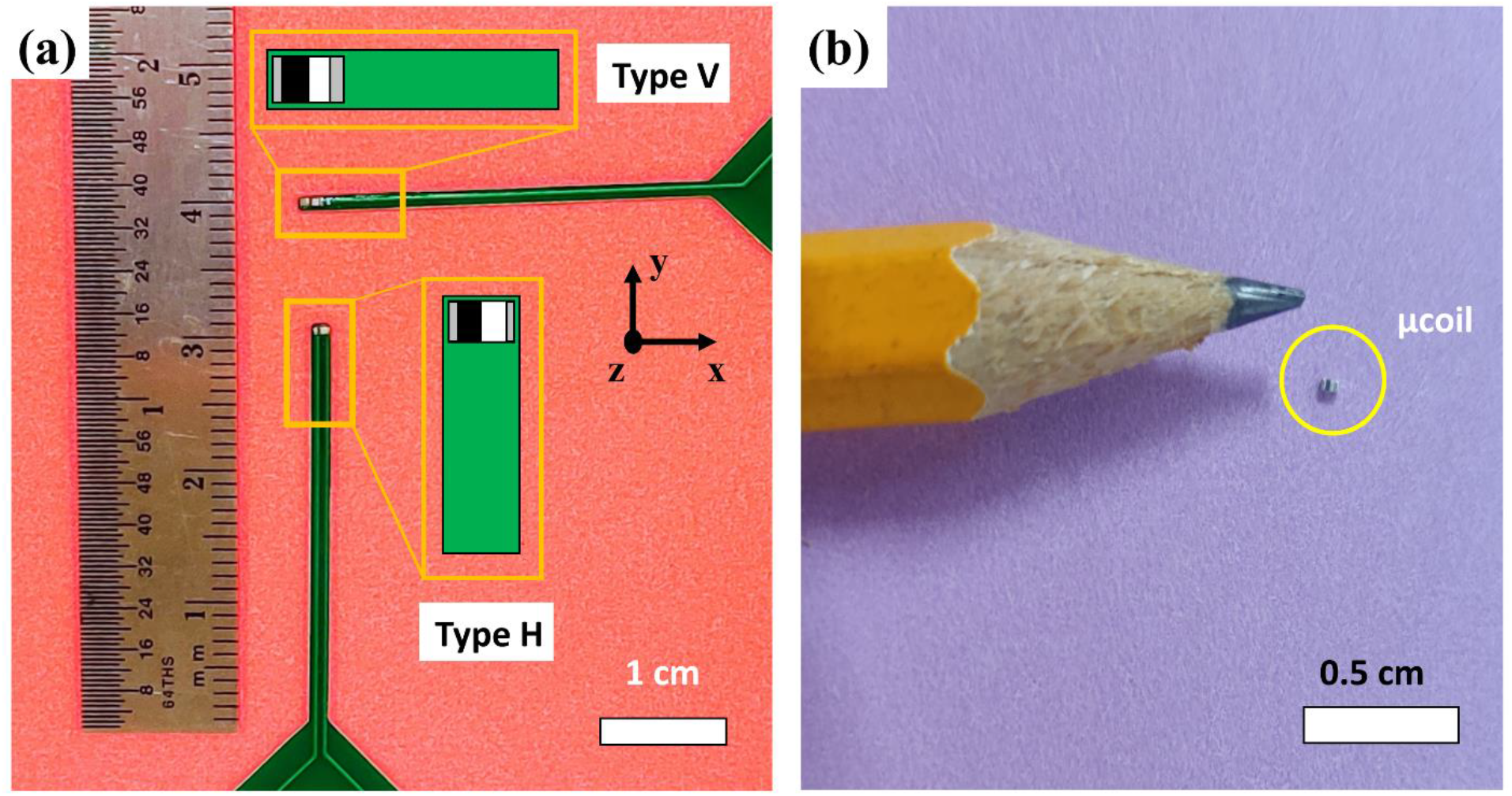
(a) The <u>Mag</u>netic <u>Pen</u> (MagPen) prototype fabricated in 2 different orientations: Type <u>H</u>orizontal or Type H & Type <u>V</u>ertical or Type V. The implant or the magnetic microcoil (μcoil) is situated at the tip. (b) The comparison of the mm-size of the μcoil with respect to the tip of a pencil (Dixon No. 2 HB pencil).

In the study of neural activation capability of a micromagnetic implant, ensuring complete blockage of leakage current and capacitive current from the prototype is extremely essential. This can be achieved by encapsulating the μcoil with a biocompatible polymer coating. In this study, we provided a 10 μm thick Parylene-C coating to the tip of the MagPen using the SCS-Labcoater Parylene-C deposition system. An effectively sealed anti-leakage current coating for the MagPen prototypes were validated by measuring the impedance between one terminal of MagPen (see Fig. S1) and another electrode in a laboratory-made standard artificial cerebrospinal fluid (aCSF) solution. Standard aCSF used was a composition (in mM) of 124 NaCl, 2 KCl, 2 MgSO_4_, 1.25 NaH_2_PO_4_, 2 CaCl_2_, 26 NaHCO_3_, and 10 D-glucose [23]. If the measured impedance was 5 MΩ and above, the prototype passed our quality control test. As further discussed in Supplementary Information S2, limiting the thickness of this Parylene-C coating is also essential as the induced electric field drops rapidly with distance.

### 2.2. The electrical circuit characteristic of the μcoil

LCR meter (Model no. BK Precision 889B; see Table S1 in Supplementary Information S1) was used to measure the resistance (R), inductance (L) and capacitance (C) values of the μcoil implant. The μcoil reduced to a RL series circuit and as per Faraday’s Laws of Electromagnetic Induction, the electromotive force (emf), is directly proportional to the series inductance (L_s_). The expression is given by: 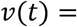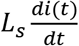, where *v*(*t*) is the emf induced in an electrical circuit (here, neurons: the biological circuits) due to a time (t)-varying magnetic field (B(t)); *v*(*t*) contributes directly to the induced electric field which is used to stimulate the neurons and 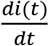 is the time derivative of the current through the μcoil (i(t)) through the inductor or μcoil. The DC resistance of the μcoil (R_DC_) contributes solely to heat dissipation from the μcoils during neuron activation [20,24,25] due to Joule heating (*heat dissipated* = *i*(*t*)^2^*R*_*DC*_*t*). Previous studies by Minusa *et al*. [26] have reported a temperature increase of ~1 °C in aCSF when the μcoils were being driven by the alternating current necessary for neural activation. With a R_DC_ value ranging between 1.8 Ω − 2.2 Ω for this μcoil, when compared to the μcoil model used in our previous work (R_DC_ = 3.8 – 4.4 Ω) [21], we expect to have the heating effect to be of 2 orders of magnitude lower.

### 2.3. Animal handling & surgery

All *in vivo* experiments were performed in accordance with a University of Minnesota approved Institutional Animal Care and Use Committee (IACUC) protocol. Seven 349±91 gram Long-Evans (L/E) rats were administered 1200 mg/kg Urethane anesthesia intraperitoneal injection and given 4 mg/kg Lidocaine and 2 mg/kg Bupivacaine local anesthesia subcutaneous injection at the incision site. Urethane was used to preserve the neurochemical environment of the animal[27]. The animal was placed on an appropriate heating device and supplemental oxygen was given via nose cone throughout the experiment. A 30 mm incision exposed the dorsal-medial right quadricep muscular interstitial space where the sciatic nerve could be accessed. One cotton tipped applicator was placed between the sciatic nerve and the quadricep for effective placement of the μcoil. Anesthetic depth and vital signs were monitored every 15 minutes throughout the experiment to ensure animal comfort and homeostasis.

### 2.4. The ‘Dose’: MagPen driving circuitry

The MagPen driving circuitry and the associated waveforms are demonstrated in Fig. 2(b). The stimulus was applied to the sciatic nerve using a function generator (Model no. RIGOL DG1022Z). The stimulus were single-cycle sinusoids; each stimulus was separated by 1 sec (see Waveform in Fig. 3). The current through the μcoil (i(t)) is represented by the equation: *i*(*t*) = *A*_1_*sin2πft*, where, the variables, *A*_*1*_ is the strength/amplitude of the current through the μcoil, *f* is the frequency of the sinusoidal current. The 1/*f* component is equivalent to the duration of the micromagnetic stimulus, a significant component of the ‘dose’. Then the current, i(t) is amplified by a class-D amplifier (Model no. Pyramid PB717X) set at a constant gain A. This time-varying current when enters a μcoil generates a time-varying magnetic field (B(t)). As per Faraday’s Law of Electromagnetic Induction, this time-varying magnetic field induces an electric field (E_ind_) which is used to activate the neural tissue. This follows directly from the equation: 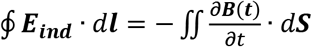. Therefore, we obtain: 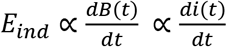. Hence, on applying a sinusoidal current waveform (i(t)) through the μcoil, we will obtain induced electric field waveform (E_ind_) in the form of time-derivate of the current. As this induced electric field activates the nerve, it will induce a sinusoidal current, *J(t)* in the nerve thereby activating axons within it (see Fig. 2(b)).

**Figure 2.**
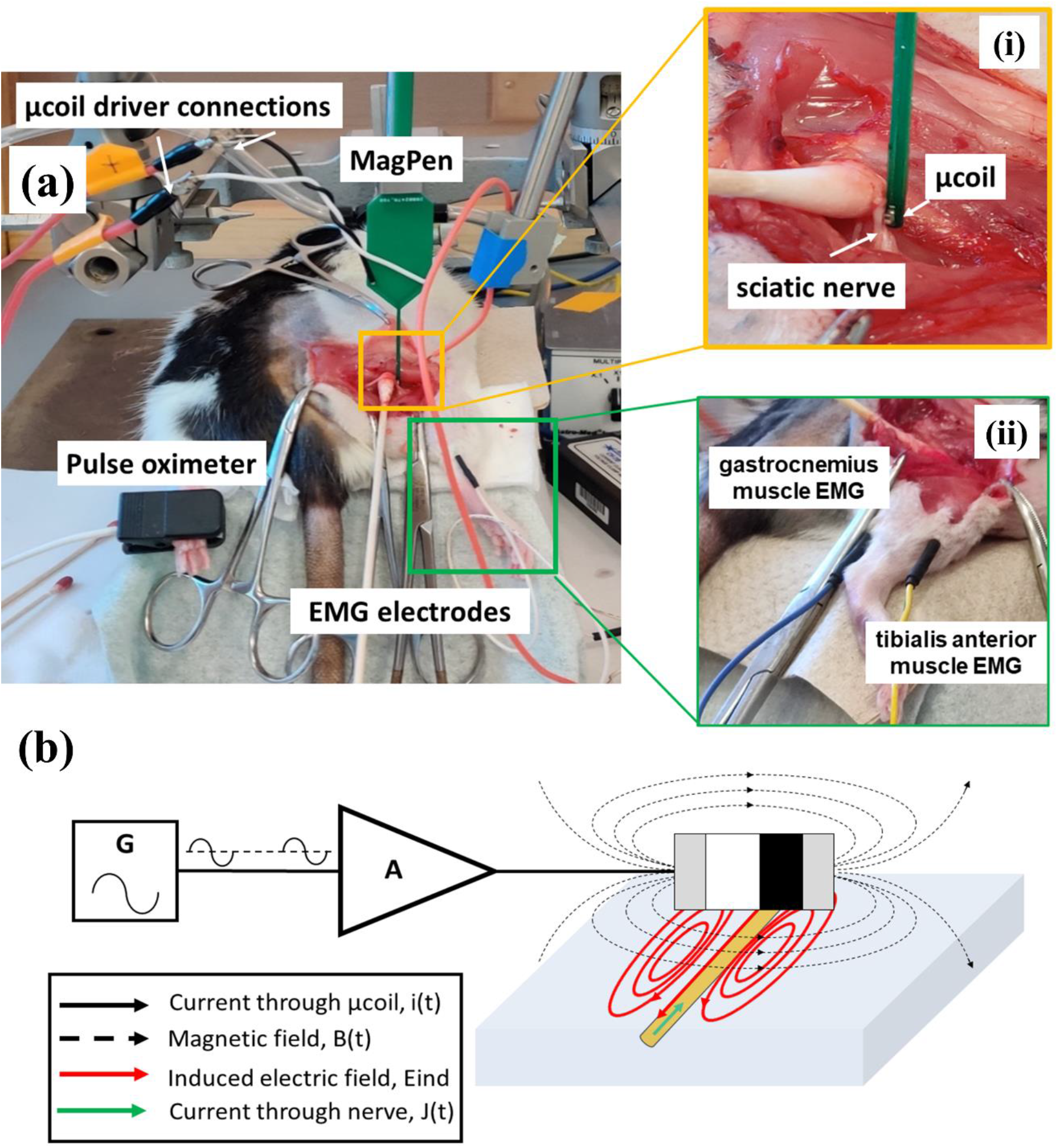
Experimental set-up for studying the effect of micromagnetic stimulation on the rat sciatic nerve. (a) The right sciatic nerve of a rat under anesthesia has been surgically exposed and the MagPen has been placed over the nerve. A pulse oximeter continually measures the heart rate and oxygen saturation in the blood of the rat. The power source connections driving the μcoil have been marked. (i) A more detailed orientation of the μcoil on the rat sciatic nerve that activated the nerve fibers. (ii) EMG electrodes have been inserted in the front and back muscles of right hind limb of the rat. (b) The MagPen driving circuit consists of a function generator (G) which generates single-cycle bursts of sinusoidal waveforms of varying amplitude and duration. This waveform is being amplified by a class-D amplifier set at a constant 10× gain thereby generating the current that drives the μcoil (denoted by solid-black lines). This amplified current, when fed into the μcoil, generates a time-varying magnetic field (denoted by dotted-black lines) which in turn induces an electric field (denoted by solid-red lines) on the nerve. This induced electric field generates a current through the nerve (denoted by solid-green lines).

**Figure 3.**
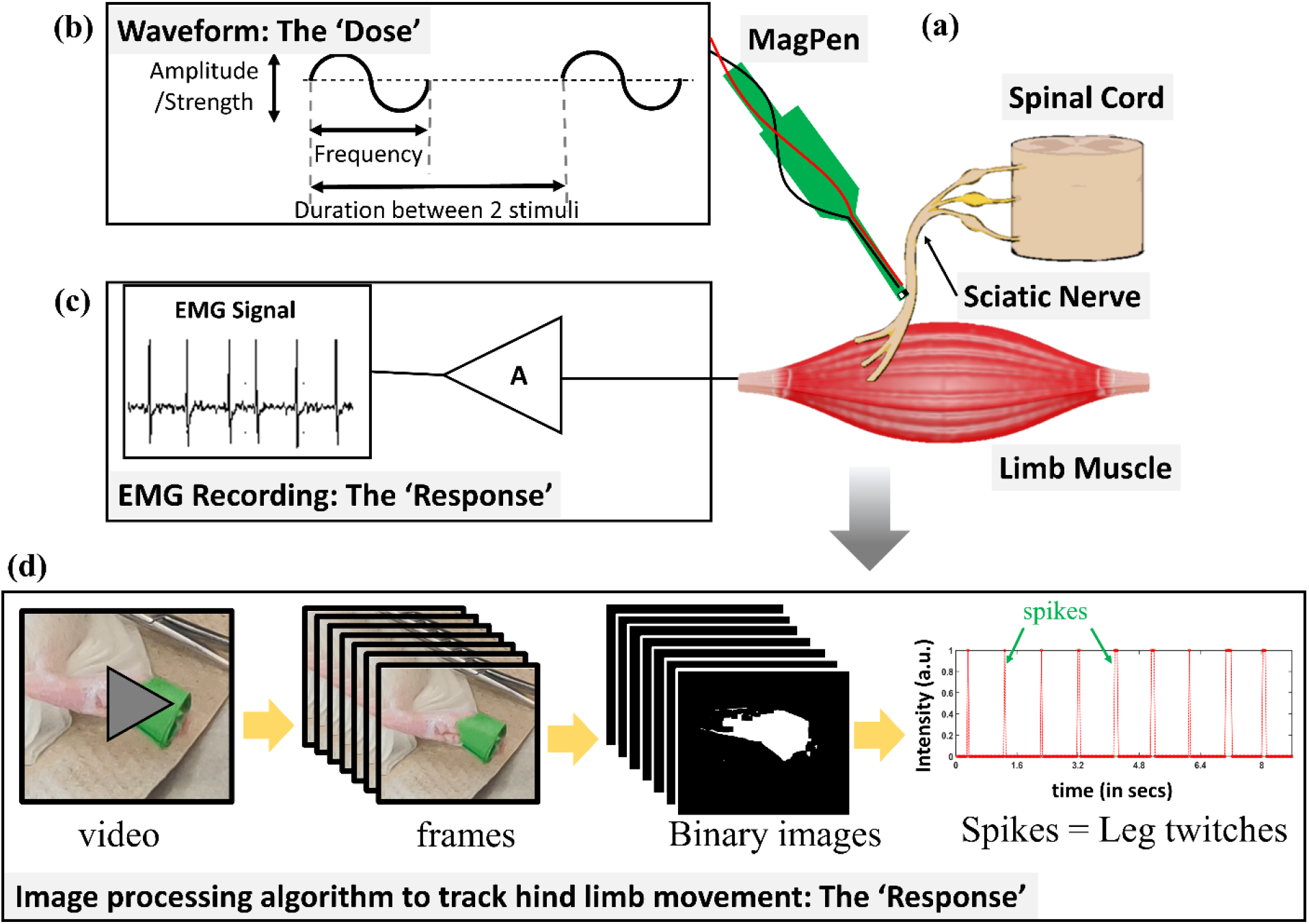
Schematic overview of the MagPen-sciatic nerve experimental set-up. (a) The sciatic nerve originates from the spinal cord and projects to the hind limb muscles. In this work, we used the MagPen (single μcoil prototype) to demonstrate micromagnetic activation of the sciatic nerve. (b) The ‘Dose’: MagPen was driven by a single wave of sinusoidal current waveform of varying amplitude and frequency; each stimulus is separated by 1 sec. (c) The ‘Response’: Stimulation of the sciatic nerve evoked electrical activity in the muscle which was recorded using electromyography (EMG) in the limb muscles. (d) In addition, the hind limb movement was tracked by applying an image processing algorithm to the video recording of the hind limb movement obtained from μMS.

### 2.5. Finite element modeling (FEM) of the μcoil implant

It is challenging to experimentally measure the magnetic field and the corresponding induced electric field from these sub-mm sized μcoils. Minusa *et al*. [26] have reported the use of miniaturized, custom-made pick-up coils for this purpose. For such measurements, it is extremely essential for the μcoil implants to be in the near vicinity of the pick-up coils. Khalifa *et al*. [28] reported the use of nitrogen vacancy (NV) magnetometers to do the same. A cost-effective yet reliable way to conduct such magnetic field and induced electric field measurements are from numerical modeling using FEM-based calculations to study the currents in neural tissues [29–31]. In this work, we used the eddy current solver of ANSYS-Maxwell [32] (ANSYS, Canonsburg, PA, United States) to study the magnetic field and the induced electric field from the μcoils. It solves an advanced form of the T-Ω formulation of the Maxwell’s equations [33]. The ceramic core μcoil dimensions, tissue slab parameters, boundary conditions and the high-resolution tetrahedral mesh size used are detailed in Table 1. All the modeling work was performed using the Minnesota Supercomputing Institute (MSI) at the University of Minnesota (8 cores of Intel Haswell E5-2680v3 CPU, 64×8=512 GB RAM and 1 Nvidia Tesla K20 GPU). The induced electric field values were then exported to be analyzed using a customized code written in MATLAB (The Mathworks, Inc., Natick, MA, USA). For the numerically simulated spatial contour plots of the magnetic field (B_x,y,z_) and induced electric field (E_x,y,z_), refer to Supplementary Information S6.

**Table 1.** Modeling parameters for the μcoil

| Parameter description | Value |
| --- | --- |
| μcoil dimension (L × W × H) | 1 mm × 600 μm × 500 μm |
| no. of turns (N) | 21 |
| Wire diameter | 7 μm |
| Tissue dimension (L × W × H) | 4 mm × 4 mm × 300 μm |
| Conductivity of tissue ($\sigma$) | 0.13 S/m |
| Air dimension (L × W × H) | 10 mm × 10 mm × 4 mm |
| Energy error (user-specified) | 1 % |
| Final solution no. of mesh elements | 410,000 |
| Adaptive passes (converged) | 6 |

### 2.6. The ‘Response’: Recording and tracking the hind limb movement activity

Stimulation of the rat sciatic nerve activates the hind limb thereby causing the leg muscle to twitch. To record the activation of the limb muscle, we used EMG electrodes (Model no. Disp. Subdermal Needle-27 ga. 12 mm from Rochester Electro-Medical Inc.) in both the tibialis anterior and gastrocnemius leg muscle of the rat (see Fig. 2(a)-ii & Fig.3(c)). EMG electrodes were connected to a headstage (Intan 32ChRHD2132) and muscle activity was recorded at 30 kHz with Open-ePhys system (https://open-ephys.org/). EMG recordings are shown in Fig S3 in Supplementary Information.

Due to the close proximity of the MagPen and the the EMG electrodes, there were significant stimulus artifacts in the EMG signals. In addition to the EMG recording, we also measured the leg twitch via video tracking and image processing. A WebCam (LogiTech C270 WebCam; 720p & 30 fps) sampled at 30 frames per second (fps) was used to record the hind limb movements. For image processing, a patch of green tape was attached to the foot. Video was collected (see Supplementary Video SV1) while stimulating the sciatic nerve with the MagPen at 1 Hz. The image data was processed using MATLAB’s Computer Vision ToolBox (The Mathworks, Inc., Natick, MA, USA) and customized code for motion tracking (see Fig. 3(d)). For data analysis, images were cropped around the green tape placed on the limb (see Supplementary Video SV2) and 9 seconds of data were analyzed. Movement was calculated by tracking changes in the green channel of the color video. A Kalman filter was used to remove noise in the resulting signal. The pixel changes were tracked & predicted inside boundary boxes (see Supplementary Information S4: Pictorial demonstration of the Image Processing Algorithm).

## 3. Results

### 3.1. Directionality of the μcoil with respect to the nerve fibers

The magnetic fields and electric fields induced by the μcoils are highly directional, so the sensitivity of the nerve to micromagnetic stimulation will be highly orientation-dependent [21,34–36]. We tested activation of the nerve with the μcoil in two orientations, horizontal (H-orientation) and vertical (V-orientation). The way the long axis of the μcoil is oriented with respect to the nerve fibers is extremely critical for successful activation of the nerve. Fig. 4(a) shows how bipolar electrical stimulation electrodes activate the nerve. Due to induction of ‘virtual cathodes’ and ‘virtual anodes’ on the nerve, it causes membrane hyperpolarization and membrane depolarization on the nerve. This phenomenon activates the nerve. For a μcoil, when the long axis of the μcoil is oriented perpendicular to a single nerve fiber (see Fig. 4(b)), due to the directionality of the induced electric fields (shown in solid-red lines), it causes similar membrane hyperpolarization and depolarization on the nerve, thereby successfully activating the nerve. For the orientations of the μcoil with respect to the nerve fibers as in Fig. 4(c) and Fig. 4(d), the directionality of the induced electric field from the μcoils, does not favor successful membrane hyperpolarization and depolarization. Our findings corroborate with the findings of Golestanirad *et al*. [34] and in Fig. 4 we have schematically represented the various orientation of the μcoil with respect to the nerve fibers (in yellow) and the directionality of the induced electric field that could potentially activate the nerve. There has not been any previous report of rat sciatic nerve activation by the μcoils, hence the correct orientation of the μcoil that would activate the nerve was unknown to us. We tried to position the μcoil with respect to the nerve fibers as in Fig. 4(b). For our experimental set-up, MagPen Type H helped us best achieve that kind of an orientation. The experimental orientation of the μcoil was found in Fig. 2(a)-i.

**Figure 4.**
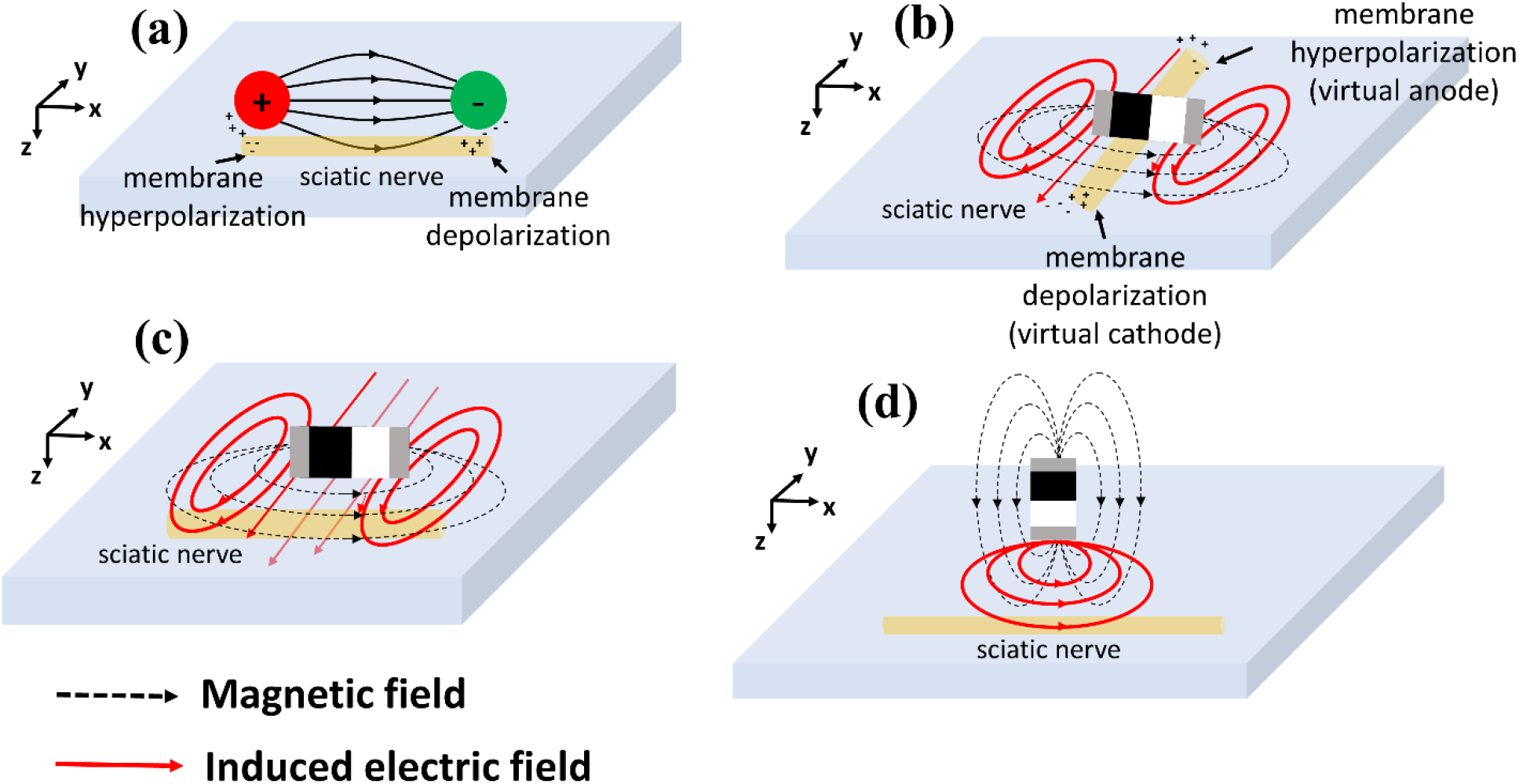
Schematic representation of importance of directionality/orientation of the μcoil with respect to the nerve for successful activation of the nerve using μMS. (a) Mechanism of activation of the sciatic nerve using bipolar electrical electrodes. (b) Desired orientation of the long axis of the μcoil perpendicular and in-plane to the nerve for activation of the nerve. (c) Long axis of the μcoil parallel to the sciatic nerve yields no activation of the nerve. (d) Long axis of the μcoil perpendicular and out-of-plane to the sciatic nerve leads to no activation of the nerve.

### 3.2. Angular significance of the μcoil with respect to the nerve tissue

Of the several interesting features of neuromodulation that μMS offers, directionality, orientation-dependence and spatial selectivity deserve special mention. Our previous report showed that MagPen has maximum chances of activating 2.672 mm^2^ (out of 16 mm^2^) of the tissue area only [21]. This spatial selectivity becomes even more precise as the distance between the μcoil and the tissue increases. However, angular orientation of the μcoil with respect to the tissue and its corresponding variation of the induced electric field has not been investigated in detail in previous studies. Here we look at the effects of angular orientation of the μcoil with respect to the tissue and its corresponding variation and induced electric fields. In this work, while trying to place the MagPen over the sciatic nerve identically, as in Fig. 4(b), there were several experimental instances (estimated to be caused by motion of the hind limb upon successful activation or breathing of the rat etc.) where the relative orientation of μcoil to the nerve changed, resulting in a change in threshold, as measured by activation of hind limb. Fig. 5 shows simulation of the effects of orientation with respect to the MagPen as predicted by the induced electric field. The induced electric field contour maps have been generated using ANSYS-Maxwell (eddy current solver) for a 4 mm × 4 mm × 300 μm biological tissue when the μcoil is being driven by a 2 A sinusoidal current of frequency 2 kHz. The distance between the μcoil and the tissue was simulated at 100 μm.

**Figure 5.**
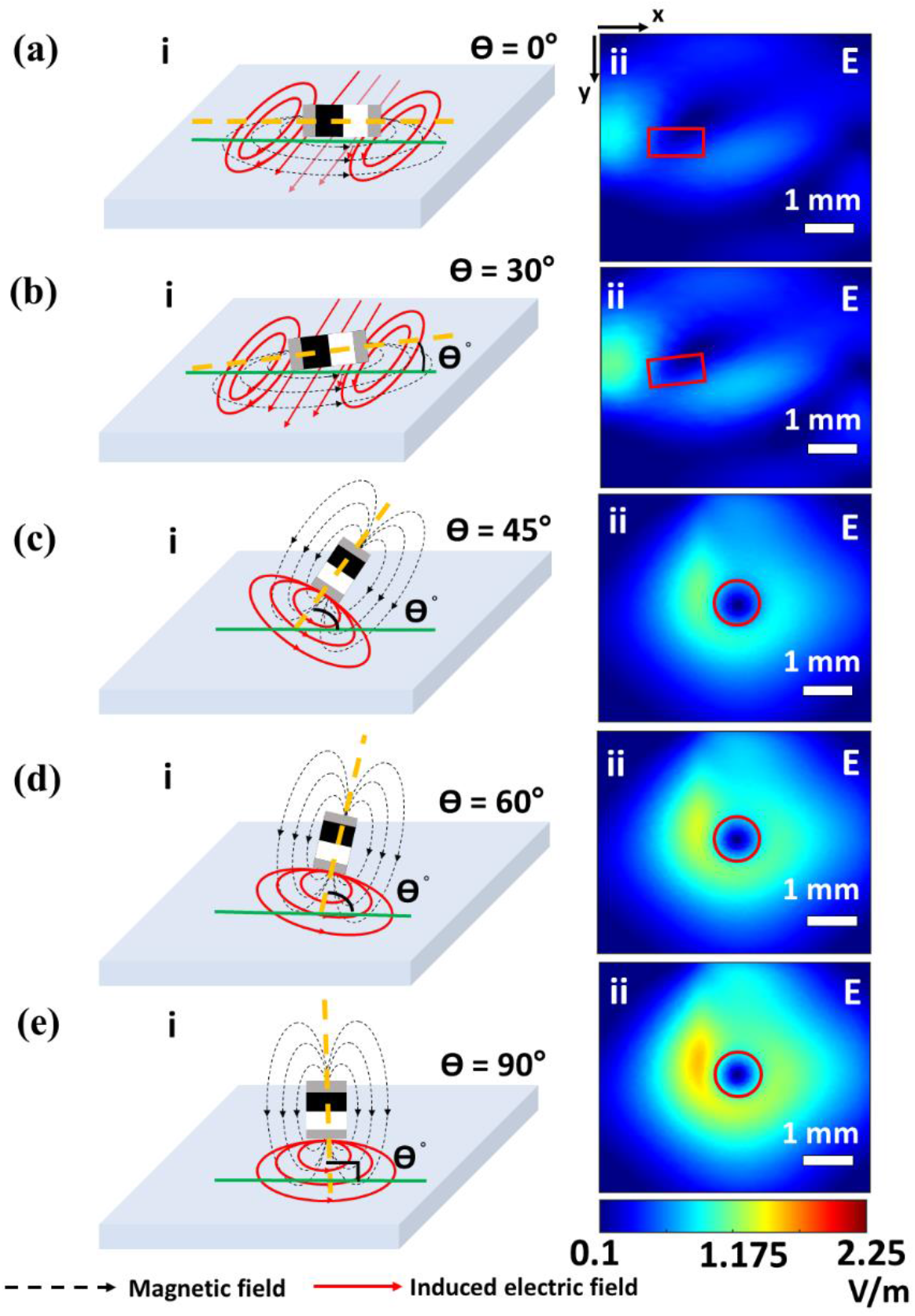
Schematic demonstration of angular dependence of μcoil with respect to the nerve orientation in the plane of the nerve tissue. Long axis of the μcoil represented by dashed-yellow lines. Orientation of the nerve fiber represented by solid-green line. Ɵ is the angle between the long axis of the μcoil and the orientation of the nerve. (a) Ɵ = 0° (i) Schematic demonstration of the long axis of the μcoil at an angle of 0֯ with respect to the nerve fiber. (ii) Numerically simulated induced electric field for the condition Ɵ = 0°. Ɵ = 30° (i) Schematic demonstration of the long axis of the μcoil at an angle of 30° with respect to the nerve fiber. (ii) Numerically simulated induced electric field for the condition Ɵ = 30°. (c) Ɵ = 45° (i) Schematic demonstration of the long axis of the μcoil at an angle of 45° with respect to the nerve fiber. (ii) Numerically simulated induced electric field for the condition Ɵ = 45°. (d) Ɵ = 60° (i) Schematic demonstration of the long axis of the μcoil at an angle of 60° with respect to the nerve fiber. (ii) Numerically simulated induced electric field for the condition Ɵ = 60° (e) Ɵ = 90° (i) Schematic demonstration of the long axis of the μcoil at an angle of 90° with respect to the nerve fiber. (ii) Numerically simulated induced electric field for the condition Ɵ = 90°. All numerical simulation for the induced electric field has been calculated for the μcoil being driven by a sinusoidal current of 2 A amplitude and 2 kHz frequency, measured on a tissue of dimension 4 *mm* × 4 *mm* × 300 μ*m*. The distance between the μcoil and the tissue was 100 μm.

### 3.3. Frequency dependence of the induced electric field

Our previous work with MagPen activating the CA3-CA1 synaptic pathway for hippocampal slices [21] also reported a numerically simulated ‘strength-frequency’ plot for μMS. When the μcoils are being driven by a time (t) -varying sinusoidal current of frequency of *f* and amplitude *A*, the current (in units of ampere (A)) through the μcoil can be expressed as:

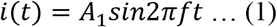

This amplitude *A*_1_ of this current through the μcoil can also be written as:

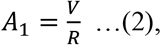

where *V* is the voltage applied to the μcoil (in units of volt (V)) from the driving circuit as in Fig. 2(b) and *R* is the resistance of the μcoil (in units of ohm (Ω)). This current *i*(*t*) generates a time-varying magnetic field *B*(*t*) given by the equation:

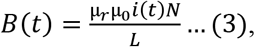

where, μ_*r*_ is the relative magnetic permeability of the medium, μ_0_ is the vacuum permeability of the medium, *N* is the number of turns in the μcoil and *L* is the length of the μcoil. This magnetic field as per Faraday’s Laws of Electromagnetic Induction induces an electric field (***E***_***ind***_) in the medium which is expressed by:

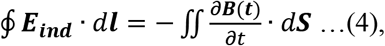

where, ***l*** and ***S*** are the contour and the surface area of the neural tissue. By substituting the expressions of ***B***(***t***) and ***i***(***t***) in equation (4), followed by some basic differentiation, we obtain the value of ***E***_***ind***_ as:

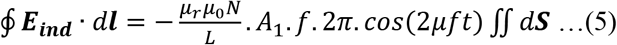

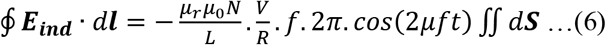

From equation (5), we obtain that *E*_*ind*_ ∝ *A*_1_. *f*, indicating the value of induced electric field is directly proportional to the two significant component of the ‘dose’ (see section 2.4) that activates the rat sciatic nerve: the amplitude of the current driving the μcoil (*A*_1_) and the frequency of the sinusoidal current (*f*). Keeping all the other variables in equation (5) as constant, to achieve a particular value of *E*_*ind*_ necessary for neural activation, if either one component of the ‘dose’ for μMS (*A*_1_ or *f*) is increased, the other component can be decreased. This frequency dependence of the induced electric field is a unique feature for μMS which follows directly from the Faraday’s Laws of Electromagnetic Induction. Fig. 6(a) & Fig. 6(b) shows the increase in magnitude of the induced electric field with increase in frequency for MagPen orientations Type V and Type H respectively; the current driving the μcoils have been kept constant. Equation (6) has been obtained from equation (5) by substituting equation (2) in equation (5). In this work showcasing μMS of the rat sciatic nerve, we have experimentally demonstrated this frequency-dependent phenomena (details explained in section 3.5). For the numerically simulated spatial contour plots of the magnetic field (B_x,y,z_) and induced electric field (E_x,y,z_), refer to Supplementary Information S6.

**Figure 6.**
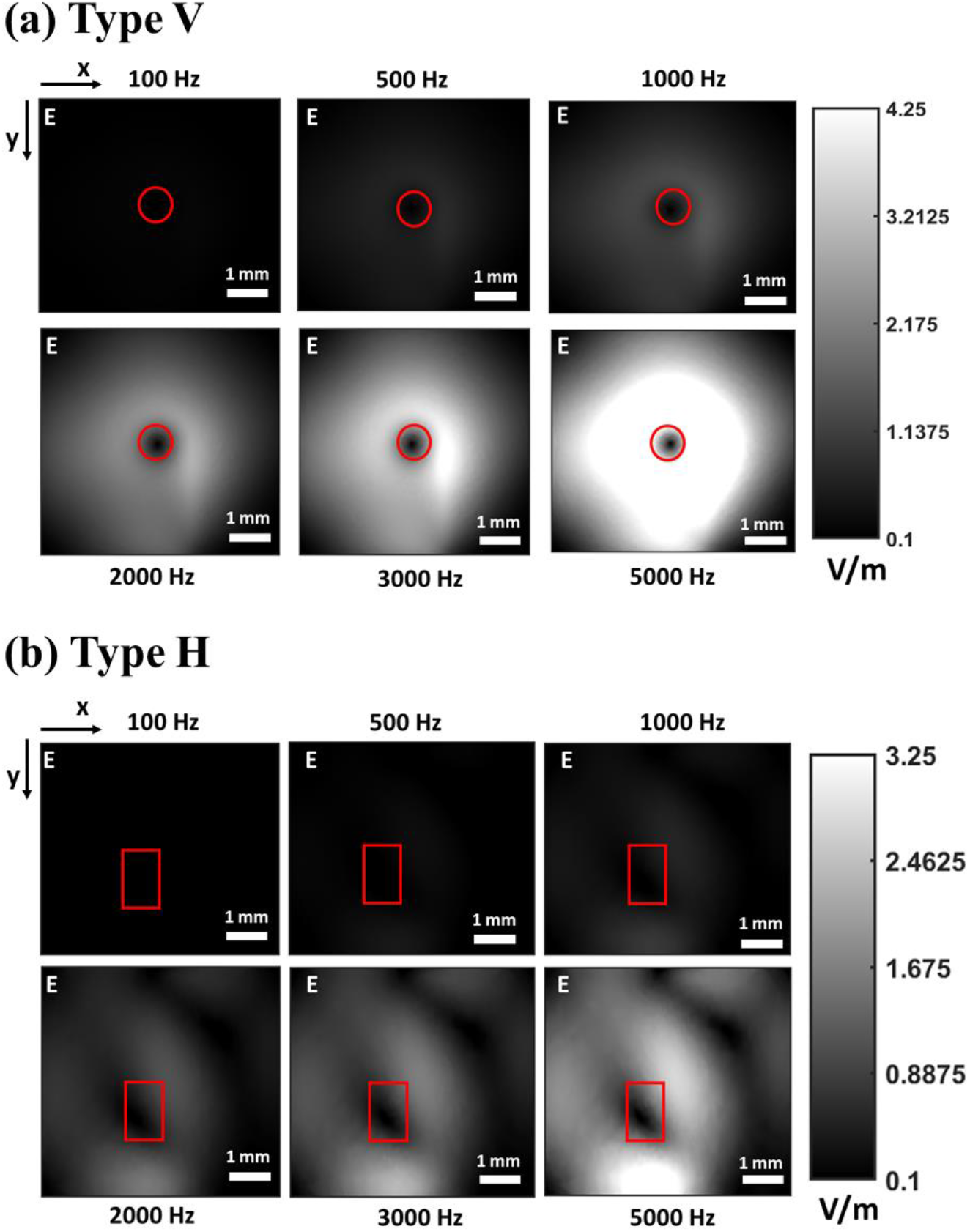
FEM modeling shows that with increase in frequency of the sinusoidal stimulus, there is increase in induced electric value. Working window shows the numerically simulated induced electric field (E in V/m) calculated for a μcoil being driven by a current of 2 A, at varying frequency, measured on a tissue of dimension 4 *mm* × 4 *mm* × 300 μ*m*. Distance between μcoil and neural tissue is taken to be 100 μm. (a) Frequency dependence of induced electric field for MagPen Type V. (b) Frequency dependence of induced electric field for MagPen Type H. The position of the μcoil has been marked in solid-red boundary.

### 3.4. Tracking the hind limb movement using image processing algorithms

In this work, we have shown that μcoils can successfully activate the rat sciatic nerve. The successful activation of the sciatic nerve has been determined by twitches causing foot dorsiflexon. We recorded the dorsiflexion movements using videos. For image processing analysis, green tape was wrapped around the toe to facilitate image processing. Fig. 7(a) shows the green tape wrapped around the toe of the right hind limb of the rat with the MagPen held over the sciatic nerve. Fig. 7(b)-i shows the successful tracking of the green tape wrapped hind limb inside the boundary box (bbox) (marked in yellow outline in Fig. 7(b)-i & Fig. 7(b)-ii). By the image processing algorithm explained in detail in section 2.6, we successfully created a binary image of the tracked limb in Fig. 7(b)-ii. Fig. 7(b)-i is a single frame from the 9-sec video (see Supplementary Video SV2) and this tracking and conversion into binary images (as in Fig. 7(b)-ii) has been repeated over all frames of the video. Fig. 7(c) is the sum of the temporal differences between frames was used to estimate movement of the leg. The MagPen was driving a single-cycle sinusoidal waveform of 3 V_p-p_ amplitude at a frequency of 1 kHz; stimuli were delivered at 1 sec interval. Therefore, following from equation (1) & (2): *V* = 3 *V*_*p*−*p*_ & *f* = 1 *kHz*. Since the micromagnetic stimulus was applied at 1 sec intervals, we must observe a leg twitch every 1 sec. Upon adding a threshold above the pixel noise (the dotted-yellow line in Fig. 7(c)), we obtained 9 distinct spikes separated by 1 sec intervals in Fig. 7(d). Those 9 spikes were observed throughout the time the MagPen was on. In this way, we have successfully tracked the hind limb movement of the rat upon μMS of the rat sciatic nerve using image processing algorithm. EMG recordings were also used to monitor muscle activation (see Supplementary Information S3). However, due to close proximity of the MagPen to the EMG electrodes, large stimulus artifacts were seen. More image processing results are provided in Supplementary Information S5. There we have applied image processing algorithms on Supplementary Video SV3 and showed that the algorithm can successfully track limb movements identifying leg kicks at 3 second intervals, corresponding to the magnetic stimulation.

**Figure 7.**
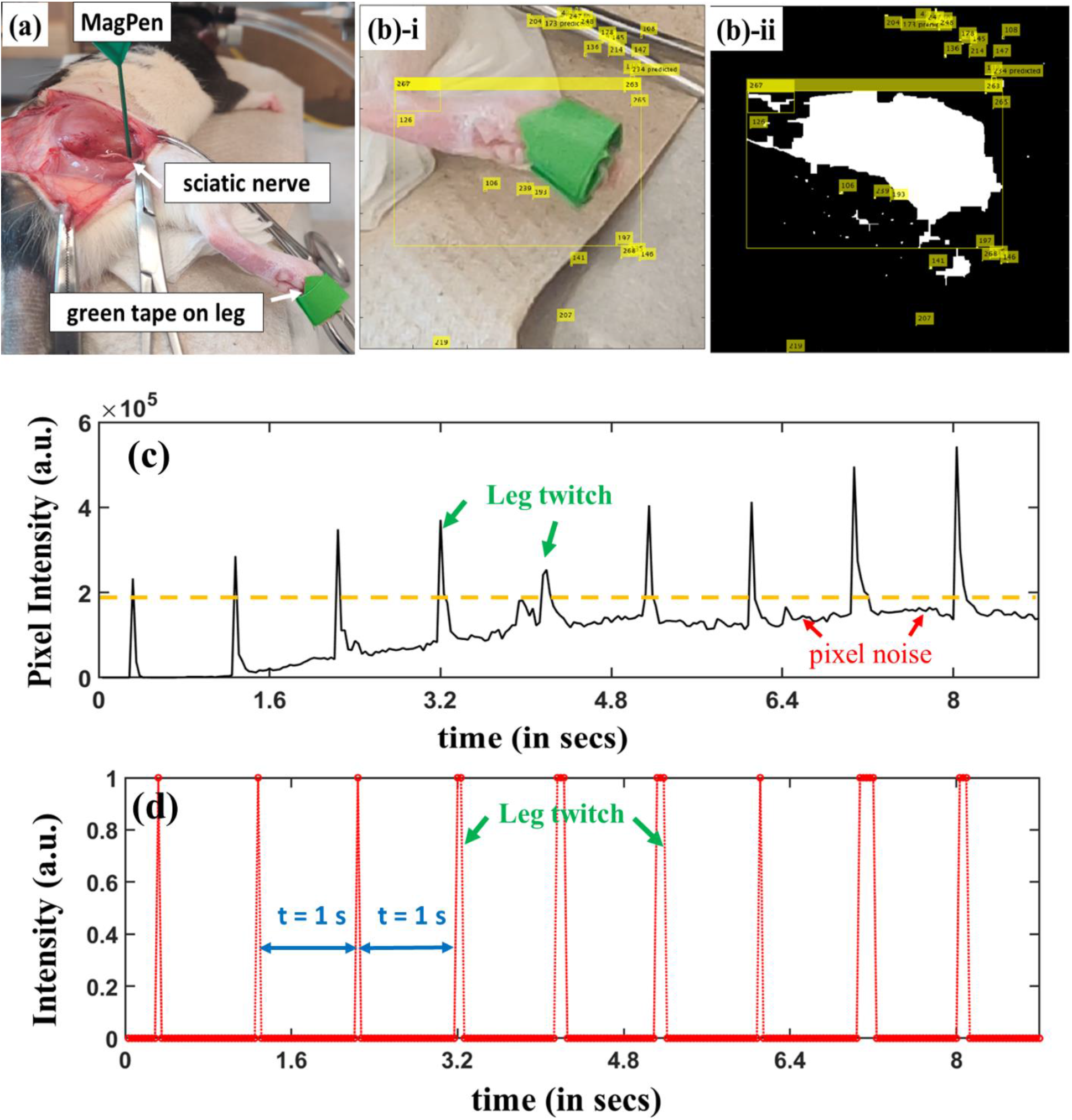
Image processing algorithm to detect leg muscle twitches/movement from a 9 sec long video sampled at 30 Hz with MagPen activating the sciatic nerve at 1 second interval. (a) A rat with right sciatic nerve surgically exposed and green tape attached to the leg. (b)-i. Applying image processing algorithm on a single frame of the 9 sec video. The algorithm creates boundary box (bbox) around rat leg keeping the green tape on the leg as a centroid for the bbox. (b)-ii. The binary image formed from the image in (b)-i, termed as ‘mask’. In this binary image, the portion that needs to be detected, *i.e*. the leg in green tape has been color-coded in white (the area under the bbox). The remaining portion of the image has been color coded in black. (c) The 9 sec video has been divided into 275 frames. The image processing algorithm was applied to each frame and reduced to the binary images as in (b)-ii. Between any 2 consecutive frames, if the leg muscle twitched, it caused the bbox to detect white patches around the green tape on the leg. Sum of the white pixels in each frame (pixel noise unfiltered). (d) To remove the pixel artefact, and other miscellaneous pixel noise from frames, a threshold was applied (dotted-yellow line in (b)-i) to the difference images. This gave rise to distinct peaks separated by 1 second, each peak corresponding to sciatic nerve activation by the MagPen. The ‘dose’ driving the MagPen was a single-cycle sinusoidal waveform of 3 V_p-p_ amplitude at a frequency of 1 kHz; each cycle separated by 1 sec. The MagPen was used to stimulate at 1 sec intervals throughout the 9 second video.

### 3.5. The dose-response relationship for μMS

Next, we varied the ‘dose’ of the micromagnetic stimuli and measured the ‘response’, as determined by whether there was a measurable twitch in the hindlimb or not (see Fig. 8(a)). Twitch response was measured as we varied the stimulation peak-to-peak voltage (V_p-p_) and the frequency of the single sinusoidal current (corresponding to the duration of the micromagnetic stimuli). We found 4 behaviors that were later used to classify the responses: (1) no leg movement; (2) weak leg movement; (3) moderately strong leg movement; and (4) strong leg movement. Here, ‘leg movement’ refers to the right hind limb movement in the rat. The dose-response relationship reported in Fig. 8(a) were repeated with 7 rats (*n* = 7).

**Figure 8.**
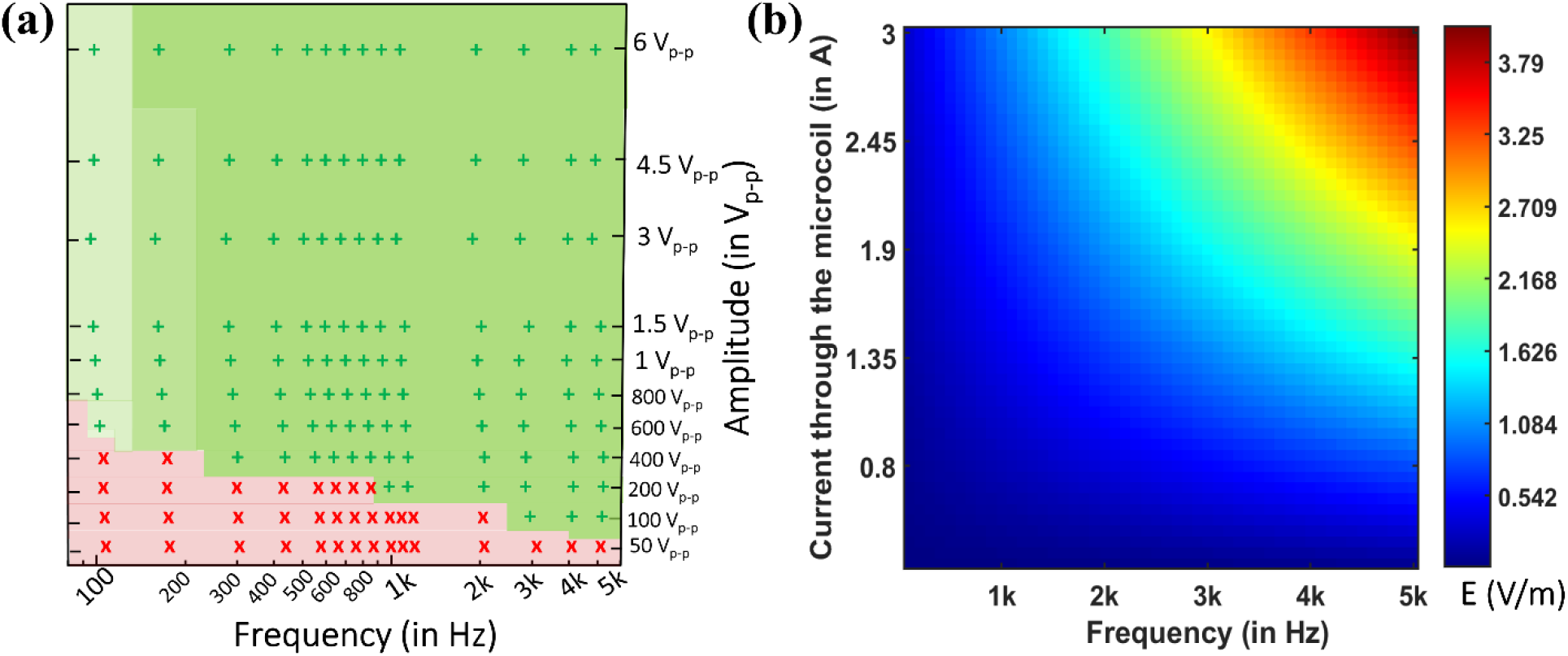
(a) The dose-response relationship experimentally measured by μMS of the rat sciatic nerve (*n*=7). In this plot, the ‘dose’ is the amplitude (in V_p-p_) and frequency (in Hz) of the μMS and the ‘response’ is the intensity of the rat hind limb muscle twitch observed. The ‘green +’ symbols indicate the μMS that elicited hind limb movements and the ‘red-x’ symbols indicate those that did not. Areas shaded in light, moderate and dark green represent parameters with weak, moderate and strong leg response, respectively. (b) The numerically simulated variation of the induced electric field (E in V/m) with the current through the μcoil (in A). 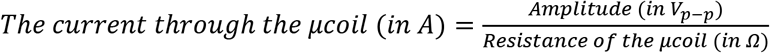.

Summarizing Fig. 8(a), we observe that at lower frequency (f) and lower voltage (V_p-p_) of the dose, there is no leg movement (see Supplementary Video SV5). At lower frequency (f) but high voltage (V_p-p_) of the dose, there is weak leg movement (see Supplementary Video SV4). At moderate frequency (f) and higher voltage (V_p-p_) there is moderately strong leg movement – (see Supplementary Video SV6). Finally, at high frequency (f) and higher voltage (V_p-p_) there is strong leg movement (see Supplementary Video SV2). Intensity of the hind limb movement is characterized in Fig. S3 (see Supplementary Information S3). This phenomenon can be explained directly from Faraday’s Laws of Electromagnetic Induction and its frequency-dependent operation, which has been better explained in section 3.3 and in Fig. 8(b). The amplitude of the response corresponds well with the amplitude of the estimated induced electric fields (from equation (6)).

## 4. Discussion

This work is the first report of micromagnetic activation of the rat sciatic nerve using sub-mm sized μcoils. We observed a dose-response relationship for μMS that follows directly from Faraday’s Laws of Electromagnetic Induction [21]. The ‘dose’ consisted of the micromagnetic stimuli where the amplitude and the frequency of the sinusoidal current waveform (or duration of the stimuli) has been controlled by an external driving circuit (see section 2.4). The ‘response’ for right sciatic nerve activation was the dorsiflexion motion of the hind limb (see section 2.6). The limb movement was quantified using video analysis. The image analysis was done using custom code developed using Computer Vision Toolbox on MATLAB and applied on the videos to track the limb movement (see section 2.6). To prevent activation of the sciatic nerve through leakage currents at the μcoils connections of the MagPen, a water-tight and biocompatible coating was used to insulate the MagPen. The thickness of the coating was less than 100 μm, so as not to affect the induced electric field because the field strength decreases rapidly with distance (see Supplementary Information S2).

The orientation of the nerve to the μcoil strongly modulated the response, as was expected by the induced electric field and justified in section 3.1. The angular dependence of the μcoil with respect to the, the nerve, and the magnitude of the induced electric field has been explained in detail in section 3.2. The dose-reponse relationship was repeated over 7 animals. Interestingly, in two animals we observed different muscle activation depending on the waveform used. In these animals, low frequency (high duration) micromagnetic stimuli induced dorsi flexion of the foot (see Supplementary Video SV7) while high frequency (low duration) micromagnetic stimuli generated a knee flexion (see Supplementary Video SV8). Most frequencies on the dose-response relationship in Fig. 8(a) generated knee flexion. The green tape was attached to the toe of the rat hind limb for even more precise RGB pixel tracking of the limb. All analysis was based on video processing, but was also confirmed with EMG recordings from the anterior and posterior muscles of the hind limb which has been reported in Supplementary Information S3.

## 5. Conclusions

This experimental work reports the activation of the rat right sciatic nerve using sub-mm sized μcoils by tracking the right hind limb motion. For the first time we experimentally report the dose-response relationship for micromagnetic stimuli using the MagPen prototype. The orientation dependence of the nerve with respect to the μcoils was measured. The resulting relationship between orientation and stimulus waveform was predicted directly from the Faraday’s Laws of Electromagnetic Induction.While μMS resulted in similar responses as electrical stimualtion, there are important differences. First, the μcoils do not have a electrochemical interface, so they may not be as sensitive to biofouling as electrodes. Second, magnetic fields fall off faster than electric fields, so μcoils may have much higher specificity than monopolar, or even bipolar electrodes. Three, MagPen induces electric fields in the tissue, which increases with frequency for the same amplitude driving the coil.This work demonstrates that micromagnetic stimulation using sub-mm sized μcoils can be used as an alternative to electrode technolgy for focal neural stimualtion. Finally, these μcoils have advantages over electrode technolgy because there is no electrochemical interface to the tissue that can degrade with use or biofouling and the spatial selectivity is much higher.

## Supplementary Videos

**SV1**: The ‘dose’ - single-cycle burst of sinusoidal waveform of 3 V_p-p_ amplitude at a frequency of 1 kHz; each burst separated by 1 sec. (Complete video)

**SV2**: The ‘dose’ - single-cycle burst of sinusoidal waveform of 3 V_p-p_ amplitude at a frequency of 1 kHz; each burst separated by 1 sec. (9 sec, zoomed-in video used in image processing)

**SV3**: The ‘dose’ - single-cycle burst of sinusoidal waveform of 3 V_p-p_ amplitude at a frequency of 1 kHz; each burst separated by 3 sec. (15 sec, zoomed-in video used in image processing)

**SV4**: The ‘dose’ - single-cycle burst of sinusoidal waveform of 3 V_p-p_ amplitude at a frequency of 100 Hz; each burst separated by 1 sec.

**SV5**: The ‘dose’ - single-cycle burst of sinusoidal waveform of 100 mV_p-p_ amplitude at a frequency of 100 Hz; each burst separated by 1 sec.

**SV6**: The ‘dose’ - single-cycle burst of sinusoidal waveform of 4.5 V_p-p amplitude_ at a frequency of 200 Hz; each burst separated by 1 sec.

**SV7**: Toe-lift Motion – The ‘dose’ - single-cycle burst of sinusoidal waveform of 4.5 V_p-p_ amplitude at a frequency of 100 Hz; each burst separated by 1 sec.

**SV8**: Pullback Motion – The ‘dose’ - single-cycle burst of sinusoidal waveform of 4.5 V_p-p_ amplitude at a frequency of 2 kHz; each burst separated by 1 sec.

## Notes

The authors declare no conflict of interest.

## Acknowledgements

This study was financially supported by the Minnesota Partnership for Biotechnology and Medical Genomics under award number ML2020. Chap 64. Art I, Sec11on 4. R.S. acknowledge the 3-year College of Science and Engineering (CSE) Fellowship awarded by University of Minnesota, Twin Cities. Research reported in this publication was supported by the University of Minnesota’s MnDRIVE (Minnesota’s Discovery, Research and Innovation Economy) initiative. The authors would also like to thank useful discussions Dr. Winfried A. Raabe, M.D. from the Department of Neurosurgery; Kendall H. Lee, M.D., PhD, Charles D. Blaha, PhD and Yoonbae Oh, PhD from Mayo Clinic, Rochester, MN. Portions of this work were conducted in the Minnesota Nano Center (MNC), which is supported by the National Science Foundation through the National Nano Coordinated Infrastructure Network (NNCI) under Award Number ECCS-2025124. J.P.W and R.S. also thank the Robert Hartmann Endowed chair support.

