## Supplementary Information for "Micromagnetic Stimulation (μMS) Dose-Response of the Rat Sciatic Nerve"

### S1. RLC measurements using LCR meter from the µcoil

LCR meter (Model no. BK Precision 889B) was used to measure the Resistance (R), inductance (L) and capacitance (C) of the µcoil at 1 kHz frequency (see Table S1). The electrical equivalent of the µcoil reduces to that of a series RL circuit for reasons explained explicitly in [1]. The parasitic value of the µcoil may vary depending on the way the µcoils have been soldered to the electrical pads of the PCB. The complete prototype of the MagPen is shown in Figure S1.

In this work, the µcoil with Model no.: TDK Corporation MLG1005SR10JTD25 has been used. Due to component shortage, if this particular model goes out of stock, one may look for other µcoils with similar electrical characteristics, i.e. one that reduces to series RL circuit, probably with even lower DC resistance to further reduce the Joule heating from the µcoils.

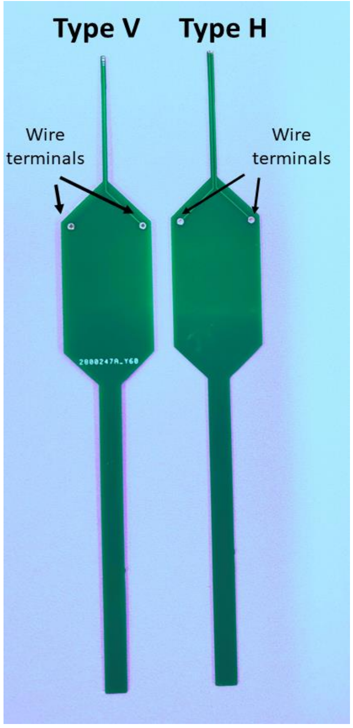


Table S1. RLC measurements of the µcoil

| Parameters | Value |
|---|---|
| Type of coil | solenoid |
| No. of turns (N) | 21 |
| Resistance R | 1.8 – 2.3 Ω |
| Series Inductance $L_s$@1kHz | 0.654 – 0.71 µH |
| Series Capacitance $C_s$@1kHz | 25 mF |
| Parallel Inductance $L_p$@1kHz | 4 mH |
| Parallel Capacitance $C_p$@1kHz | 76.83 nF |

**S2. Variation of the induced electric field with increase in distance (d) between nerve & µcoil**

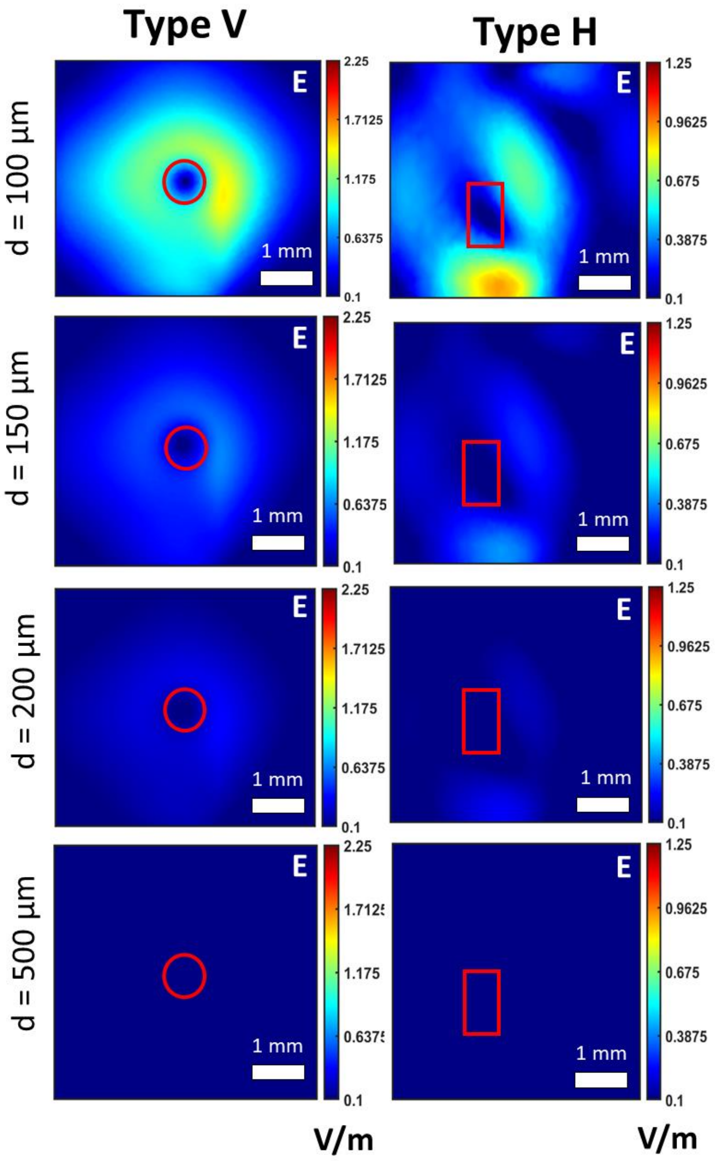


Figure S2. The rapidly decreasing induced electric field with increase in distance (d) between the nerve and the µcoil for both conditions, MagPen Type V and Type H. The outline of the µcoil highlighted in red for both Type V and Type H orientations.

The reason why controlling the precise thickness of the anti-leakage current coating, Parylene-C, is so important in this work is because the induced electric field (E) from these micrometer-sized coils attenuates significantly with distance d between the µcoils and the nerve. For both orientations of the MagPen, Type V and Type H, Fig S2 shows the rapid attenuation of the induced electric field from distance d = 100 µm to d = 500 µm. All data were simulated on ANSYS-Maxwell (eddy current solver) for a tissue of size 4 mm × 4 mm × 300 µm and a µcoil driven by a 2 kHz sinusoidal current of 2 A measured at varying distances (d) between the µcoil and the nerve.

### S3. Recording of muscle activity from the rat hind limb using EMG

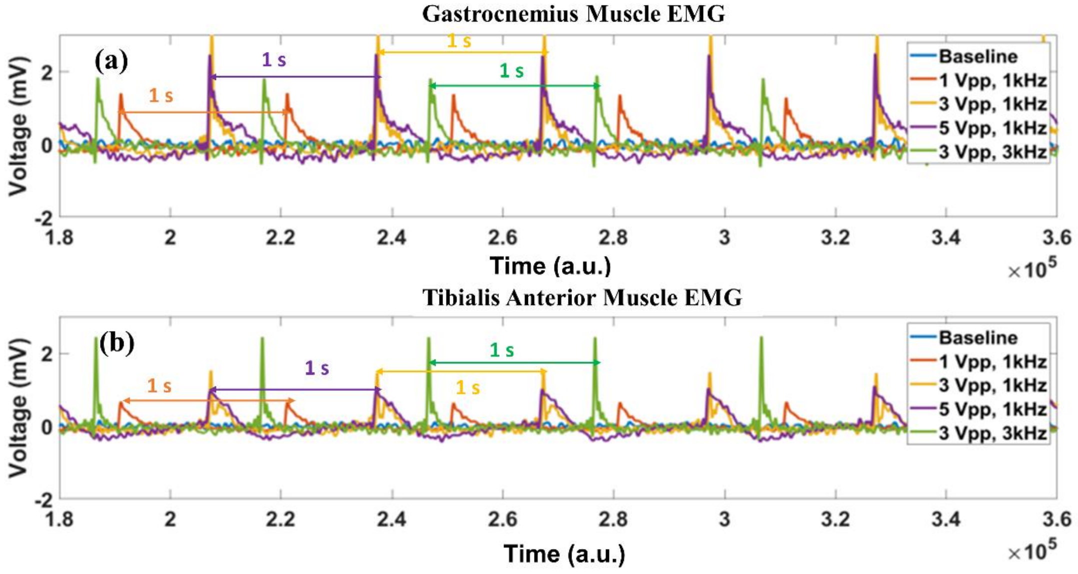


Figure S3. EMG recordings from (a) Posterior muscle of the rat hind limb. (b) Anterior muscle of the rat hind limb.

Fig. S3 compares the posterior (see Fig. S3(a)) and anterior muscle (see Fig. S3(b)) EMG recordings for the rat hind limb for varying 'dose' of the micromagnetic stimuli. The details about the EMG recording have been made in section 2.6. In Fig. S3, the solid-blue line represents the baseline signal while the orange, yellow, purple and green lines represent the micromagnetic stimuli dose at $(V = 1\, V_{p-p}, f = 1\, kHz)$, $(V = 3\, V_{p-p}, f = 1\, kHz)$, $(V = 5\, V_{p-p}, f = 1\, kHz)$ and $(V = 3\, V_{p-p}, f = 3\, kHz)$, respectively (see equation (6)). From Fig. S3(a), we see that for a constant frequency at 1 kHz, as we increase the amplitude, we get a stronger response. Whereas, if we keep the amplitude constant at 3 Vp-p but increase the frequency from 1 kHz to 3 kHz, a stronger response is recorded from EMG at 3 kHz. This follows directly from the dose-response relationship supported by Faraday's Laws of Electromagnetic Induction explained in Fig. 8 (see section 3.5). A similar observation has been supported by the EMG recordings from the anterior muscle. Most of the motion for the hind limb observed for these doses for the micromagnetic stimuli are a dorsiflexion motion (see section 4).

To superimpose recordings to compare the amplitudes at different doses, the time scale has not been calculated correctly by including the sampling rate in Fig. S3. However, the time duration difference between two consecutive spikes is 1 sec. The sampling rate for these EMG recordings were 30,000 samples per second.

### S4. Pictorial Demonstration of the Image Processing Algorithm

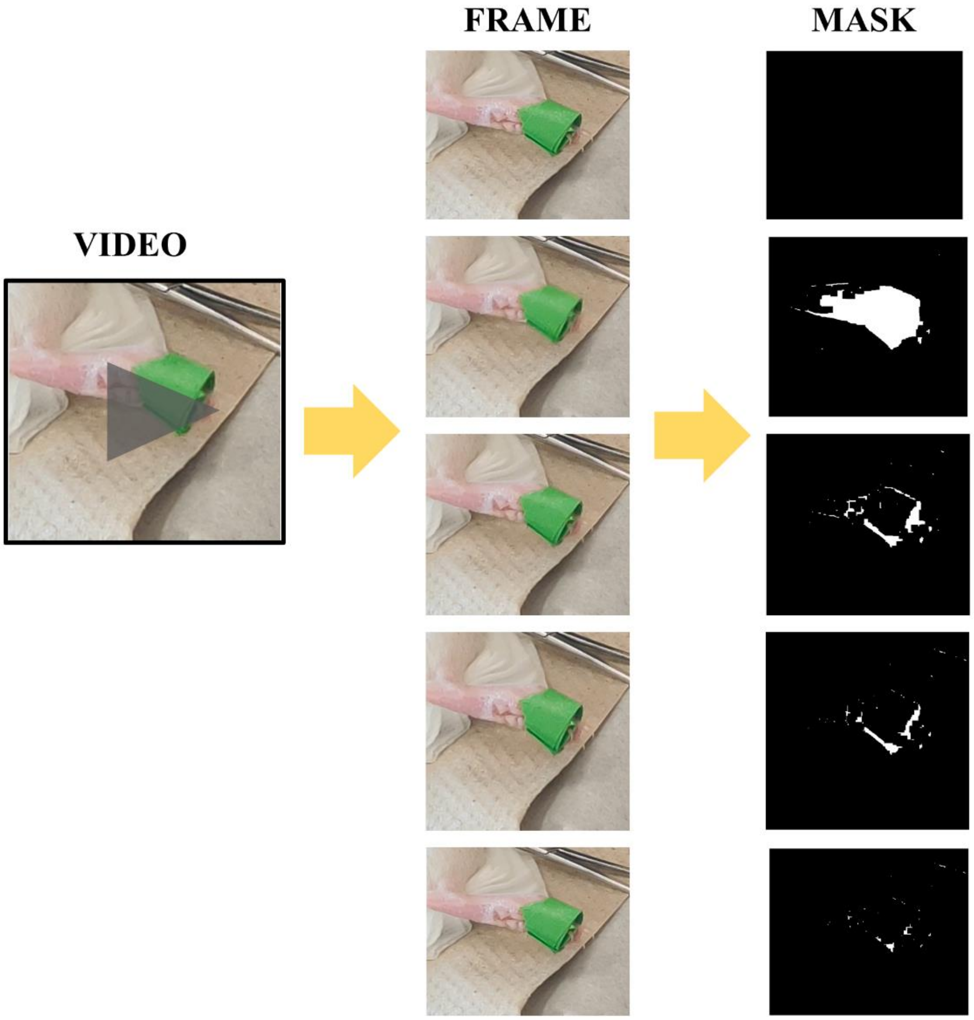


Figure S4. Pictorial demonstration of the image processing algorithm. The video for the limb movement has been sub-divided into several frames (Only 5 consecutive frames are shown here). The green tape has been tracked in the algorithm in the form of white patches only when the limb moves. Each frame has been converted into a binary mask (black and white images) which can be read as a 2D matrix of 1's (white) and 0's (black).

Using the Computer Vision Toolbox on MATLAB, we have developed this Image Processing Algorithm whose steps have been pictorially demonstrated in Fig. S4. Supplementary Video SV2 has been used for the demonstration. The 9 second video has been divided into 275 frames (1 second corresponds to 30 frames approximately) and hence the algorithm generated 275 masks. For better accuracy of the algorithm, we have attached a green tape onto the foot of the rat such that the red-green-blue (RGB) pixel tracking becomes easier and more accurate. Successful demonstration of the limb tracking has been demonstrated in Supplementary Information S5.

### S5. Image Processing Algorithm Tracking Hind Limb Movement

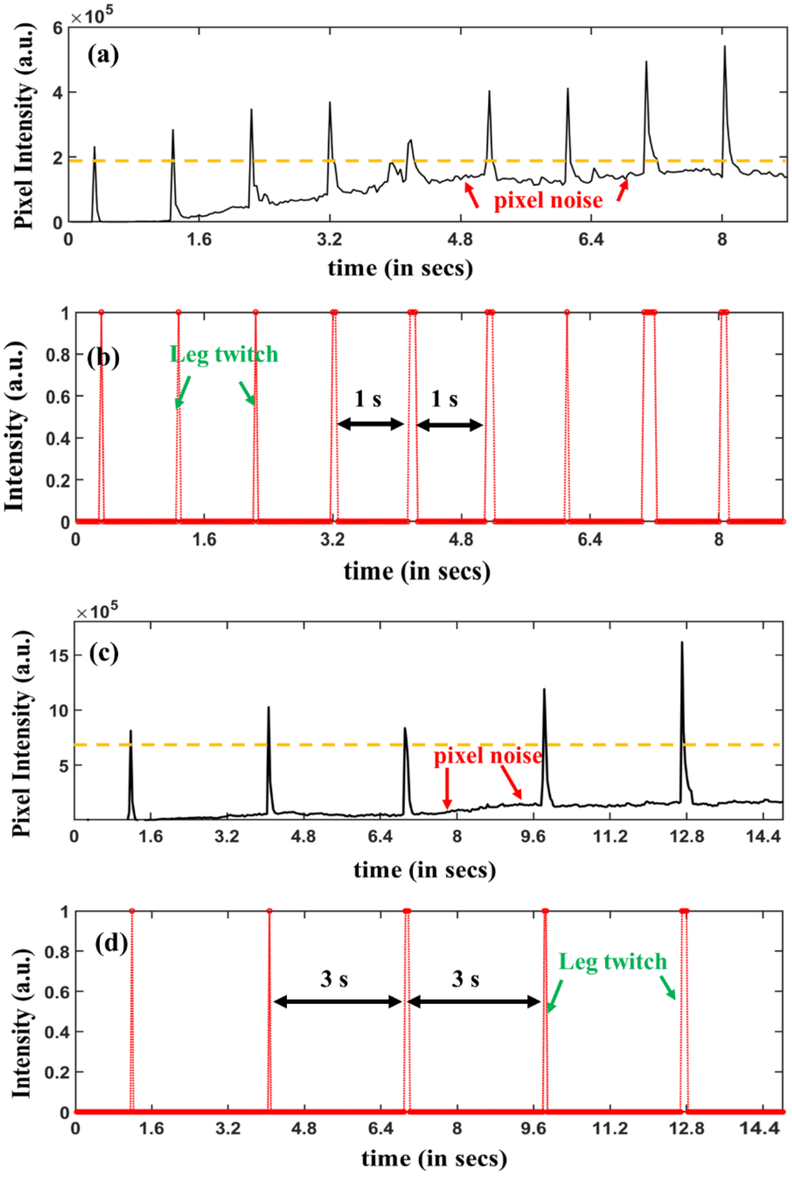


Figure S5. Tracking limb movement from Supplementary Video SV2. The micromagnetic stimuli applied to the MagPen was single-cycle sinusoidal bursts of amplitude A = 3 Vp-p, f = 1 kHz and each stimulus separated by 1 second. MagPen was on throughout the 9 second video. (a) Sum of the white patches in the binary mask image represented in raw data (including pixel noise). (b) Leg kicks represented by the spikes at every 1 second. Tracking limb movement from Supplementary Video SV3. The micromagnetic stimuli applied to the MagPen was single-cycle sinusoidal bursts of amplitude A = 3 Vp-p, f = 1 kHz and each stimulus separated by 3 seconds. MagPen was on throughout the 15 second video. (c) Sum of the white patches in the binary mask image represented in raw data (including pixel noise). (b) Leg kicks represented by the spikes at every 3 seconds.

Two different videos SV2 & SV3 showing the successful tracking of the hind limb movement using the custom designed image processing algorithm. Each of the two conditions have different micromagnetic stimulus driving the MagPen. For the cases in Fig. S5(a)&(b), the MagPen was driven by single-cycle sinusoidal bursts of amplitude A = 3 Vp-p, f = 1 kHz and each stimulus separated by 1 second. For the cases in Fig. S5(c)&(d), the MagPen was driven by single-cycle sinusoidal bursts of amplitude A = 3 Vp-p, f = 1 kHz and each stimulus separated by 3 seconds.

**S6. Spatial components of the magnetic flux density for MagPen**

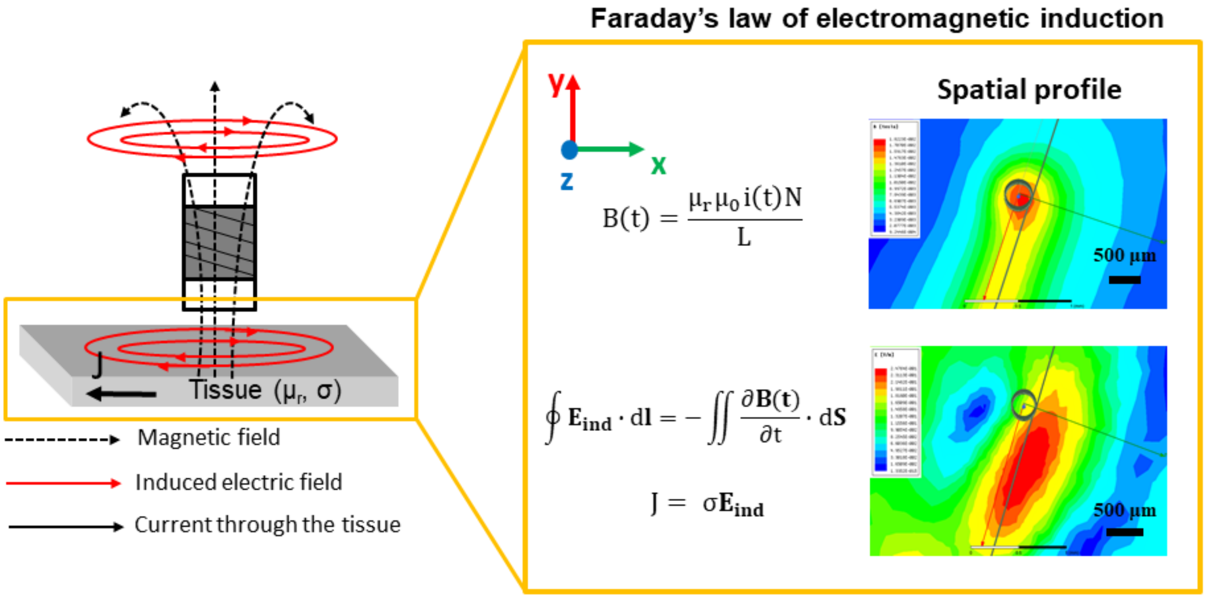


Figure S6. The spatial components of magnetic flux density, B(t) and induced electric field ($E_{ind}$) measured for the µcoil and neural tissue distance 300 µm for a sinusoidal current, i(t) of amplitude 2 A and frequency 2 kHz. Modeling performed on ANSYS-Maxwell (eddy current solver).

$\mu_r$ and $\mu_0$ are the relative permeability of the neural tissue and vacuum permeability respectively. σ is the conductivity of the neural tissue. The current that will flow through a neural tissue on stimulation by the induced electric field from these µcoils will depend on σ. When current flows through the neural tissue, the neurons start firing, i.e., it is stimulated.
